# Cyanophages exhibit rhythmic infection patterns under light-dark cycles

**DOI:** 10.1101/167650

**Authors:** Riyue Liu, Yue Chen, Rui Zhang, Yaxin Liu, Nianzhi Jiao, Qinglu Zeng

## Abstract

Most living organisms exhibit diurnal rhythms as an adaptation to the daily light-dark (diel) cycle. However, diurnal rhythms have not been found in viruses. Here, we studied the diel infection patterns of bacteriophages infecting the unicellular cyanobacteria *Prochlorococcus* and *Synechococcus*, which are the most abundant photosynthetic organisms in the oceans. With lab cultures, we found that cyanophages used three infection strategies in the dark: no adsorption, adsorption but no replication, and replication. Interestingly, the former two exhibited rhythmic infection patterns under light-dark cycles. We further showed in the South China Sea and the Western Pacific Ocean that cyanophage abundances varied rhythmically, with a peak at night. Moreover, diel transcriptional rhythms of many cyanophage genes were found in the North Pacific Subtropical Gyre, which also peaked at night. Our results suggested that cyanophage infection of *Prochlorococcus* is synchronized to the light-dark cycle, which may result in a synchronized release of dissolved organic matter to the marine food web.

---

Similar to most living organisms, cyanobacteria adapt to the daily light-dark (diel) cycle by exhibiting diurnal rhythms of gene expression, metabolism, and growth (Dunlap, 1999; Cohen and Golden, 2015). Light also influences the life cycle of viruses (cyanophages) that infect the unicellular picocyanobacteria *Prochlorococcus* and *Synechococcus* (Clokie and Mann, 2006; Ni and Zeng, 2016), which are the most abundant photosynthetic organism on earth (Partensky et al., 1999; Scanlan and West, 2002). As lytic double-stranded DNA viruses, cyanophages include three morphotypes: T4-like cyanomyoviruses, T7-like cyanopodoviruses, and cyanosiphoviruses (Suttle and Chan, 1993; Waterbury and Valois, 1993; Wilson et al., 1993; Sullivan et al., 2003). Some cyanophages show light-dependent adsorption to their host cells (Cseke and Farkas, 1979; Kao et al., 2005; Jia et al., 2010). Cyanophage replication depends on light as well, and their burst sizes were greatly reduced when host cells were infected in the light and then moved to the dark (Mackenzie and Haselkorn, 1972; Sherman, 1976; Kao et al., 2005; Lindell et al., 2005; Thompson et al., 2011; Thompson et al., 2016). Moreover, cyanophage abundance in the Indian Ocean was shown to fluctuate within a 24-h period (Clokie et al., 2006). However, cyanophages have not been shown to exhibit rhythmic infection patterns under light-dark cycles.

## Some cyanophages replicate in the dark while some cannot replicate

Before studying cyanophage infection under light-dark cycles, we first examined whether cyanophages can initiate and complete the entire infection cycle in the dark. In our cyanophage collection, we had cyanomyoviruses P-HM1, P-HM2, and P-SSM2, and cyanopodoviruses P-SSP7 and P-GSP1 (Sullivan et al., 2003; Sullivan et al., 2010). They can infect *Prochlorococcus* strains MED4 (P-HM1, P-HM2, P-SSP7, and P-GSP1) and NATL2A (P-HM1, P-HM2, and P-SSM2). We incubated cyanophages with unsynchronized host cells at a phage/host ratio of 0.1 in the dark or under continuous light, and measured phage DNA copies by quantitative PCR, which provides a ~1:1 relationship with phage particle counts (Frois-Moniz, 2014). Following a routine method for cyanophage infection kinetics (Lindell et al., 2007; Thompson et al., 2011), extracellular phage DNA copies were used to measure phage adsorption to and release from the host cells, and intracellular phage DNA copies were used to measure phage replication inside the host cells.

In the dark, cyanophages showed three infection strategies. First, P-HM2 (Figures 1A, D) and P-HM1 (similar to P-HM2) did not adsorb to their host cells in the dark, which could be due to conformational changes in host receptors or phage tail fibers that can only be induced in the light (Jia et al., 2010). Second, P-SSM2 got adsorbed to the host cells in the dark, although the adsorption was lower than that in the light (Figure 1E and Supplementary Figure 1). However, P-SSM2 did not replicate in the dark (Figure 1B), suggesting that its replication is regulated by some unknown mechanism that is sensitive to light. Third, P-SSP7 (Figures 1C, F) and P-GSP1 (similar to P-SSP7) got adsorbed to and also replicated in the host cells in the dark. Furthermore, P-SSP7 progeny was released in the dark at the same time as the cultures in the light, although the burst size seemed much smaller than that in the light (Figure 1F). This is the first time cyanophages were found to complete the entire infection cycle in the dark.

**Figure 1.**
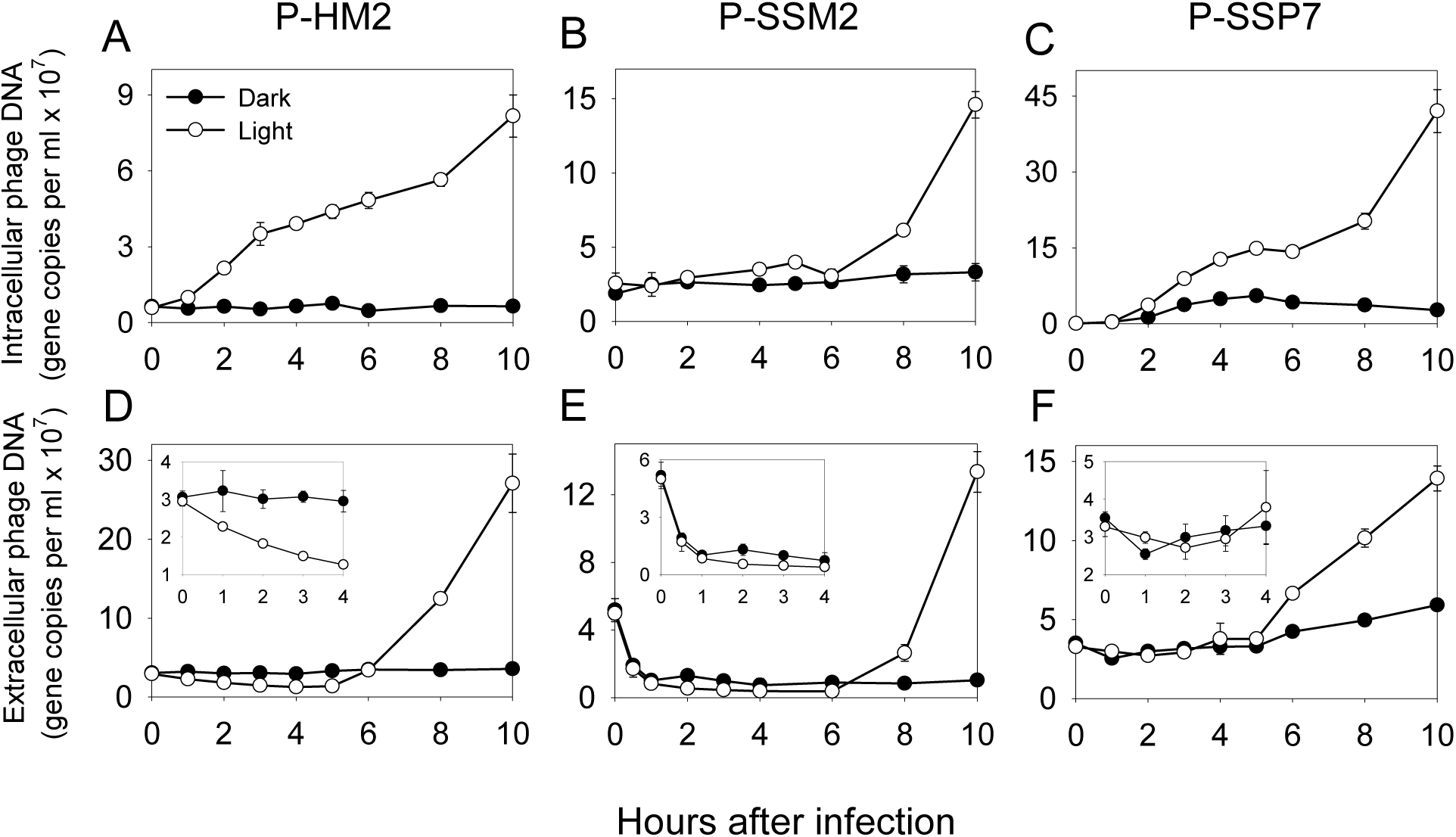
Cyanophage infection of *Prochlorococcus* in the light or in the dark. Cyanophages P-HM2 (**A** and **D**), P-SSM2 (**B** and **E**), and P-SSP7 (**C** and **F**) were used to infect their host cells at a phage/host ratio of 0.1 under continuous light or in the dark. The host for P-HM2 and P-SSP7 was *Prochlorococcus* MED4 and the host for P-SSM2 was *Prochlorococcus* NATL2A. Intracellular (**A**, **B**, and **C**) and extracellular (**D**, **E**, and **F**) phage DNA was measured by quantitative PCR. Insets in **D**, **E**, and **F** were the corresponding zoon-in data of 0–4 h. Error bars represented the standard deviations of two biological replicates and were smaller than the data point when not apparent.

## Rhythmic infection patterns of cyanophages under light-dark cycles

To compare the diel infection patterns of cyanophages P-HM2, P-SSM2, and P-SSP7, we infected synchronized *Prochlorococcus* cells grown under a 14h light:10h dark cycle (Supplementary Figure 2), where ZT (Zeitgeber Time) 0 corresponds to lights on and ZT 14 corresponds to lights off. Infections were carried out at five time points (ZT 14.5, 18 on Day 1, and ZT 0.5, 6, 12 on Day 2) (Figure 2). A low phage/host ratio of 0.02 was used to ensure that cyanophage production was not limited by the host availability for at least 2 days after the first infection (Supplementary Figure 3).

**Figure 2.**
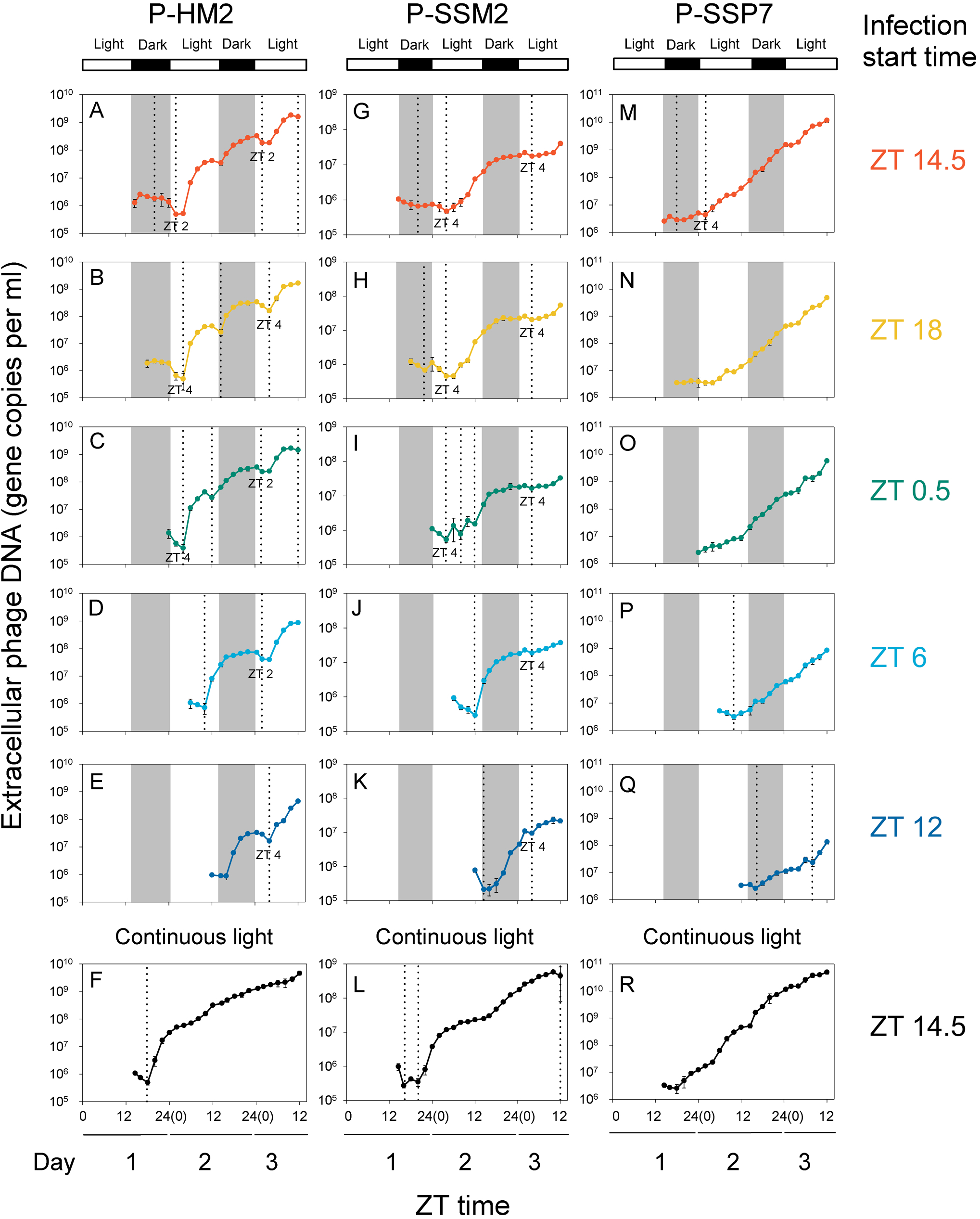
Diel infection patterns of cyanophages P-HM2, P-SSM2, and P-SSP7. At a phage/host ratio of 0.02, *Prochlorococcus* MED4 was infected by P-HM2 (**A**–**F**) or P-SSP7 (**M**–**R**) and *Prochlorococcus* NATL2A was infected by P-SSM2 (**G**–**L**). Under a 14h light:10h dark cycle (lights on at ZT 0 and off at ZT 14), synchronized *Prochlorococcus* cultures were infected at ZT 14.5 (**A**, **G**, **M**), 18 (**B**, **H**, **N**), 0.5 (**C**, **I**, **O**), 6 (**D**, **J**, **P**), and 12 (**E**, **K**, **Q**). The ZT 14.5 infections were also carried out under continuous light (**F**, **L**, **R**). Extracellular phages were measured by quantitative PCR. Error bars represented the standard deviations of two biological replicates. In each graph, the vertical dotted lines indicated the times of phage adsorption (at least 10% decrease of extracellular phage number compared to the previous time point).

When P-HM2 infections were initiated before (Figures 2A, B) or at around lights on (Figure 2C), the extracellular phage number of different infections all decreased at around ZT 2–4 on both Day 2 and Day 3. This rhythmic decrease in extracellular phage number at the same time of two days was probably due to light-dependent adsorption of P-HM2 after lights on, since this pattern was not observed when the infected cultures were moved to continuous light (Figure 2F). Moreover, when P-HM2 infections were initiated several hours after lights on (Figures 2D, E), the extracellular phage numbers also decreased at around ZT 2–4 on Day 3, indicating that P-HM2 infections initiated at different times of a day were synchronized under light-dark cycles. The rhythmic decrease in extracellular phage number was also seen in P-SSM2 infections under light-dark cycles (Figures 2G–K) but not under continuous light (Figure 2L), which could be because P-SSM2 adsorption was enhanced after lights on (Supplementary Figure 1). Besides ZT 2–4, the extracellular phage numbers of P-HM2 and P-SSM2 infections decreased at additional time points (Figures 2A–L). This might be caused by the different initiation time of each infection and/or the different latent periods of P-HM2 and P-SSM2 (Figure 1D, E). In contrast to P-HM2 and P-SSM2, P-SSP7 showed unrhythmic infection patterns both under light-dark cycles (Figures 2M–Q) and under continuous light (Figure 2R), probably because it can replicate in the dark (Figure 1F). In summary, we showed for the first time that viruses (P-HM2 and P-SSM2) exhibited rhythmic infection patterns. Since this infection rhythm can only be observed under light-dark cycles but not under continuous light, it is a diurnal rhythm, rather than a circadian rhythm, which can be maintained under continuous light.

## The light-dark cycle affects the relative fitness of cyanophages

To test whether the light-dark cycle affects the relative fitness of cyanophages, we competed two cyanophages under continuous light or at different times of a light-dark cycle (total phage/host ratio = 0.02). First, we coinfected synchronized *Prochlorococcus* cells with equal numbers of P-HM2, which showed an infection rhythm, and P-SSP7, which did not (Figure 2). P-SSP7 became the dominant phage under continuous light (Figure 3A), however, P-HM2 became the dominant phage at most times under light-dark cycles (Figures 3C–F), suggesting that P-HM2 had a fitness advantage over P-SSP7 under light-dark cycles. P-SSP7 can only defeat P-HM2 under light-dark cycles when the competition was initiated shortly after lights off (Figure 3B). This might be because P-SSP7 replicated in the dark while P-HM2 cannot (Figure 1), and after lights on P-SSP7 already had more progeny phages than P-HM2. Next, we coinfected synchronized *Prochlorococcus* cells with cyanomyoviruses P-HM2 and P-SSM2, both of which showed infection rhythms (Figure 2). We found that P-SSM2 became the dominant phage under continuous light (Figure 3G) and also at different times of a light-dark cycle (Figures 3H–L), suggesting that the light-dark cycle did not affect the relative fitness of two phages with infection rhythms. The fitness advantage of biological rhythms has been shown in cellular organisms (Johnson et al., 2017). Although we were unable to generate isogenic phages with or without an infection rhythm, our empirical evidence indicated that infection rhythms also conferred cyanophages a fitness advantage under light-dark cycles.

**Figure 3.**
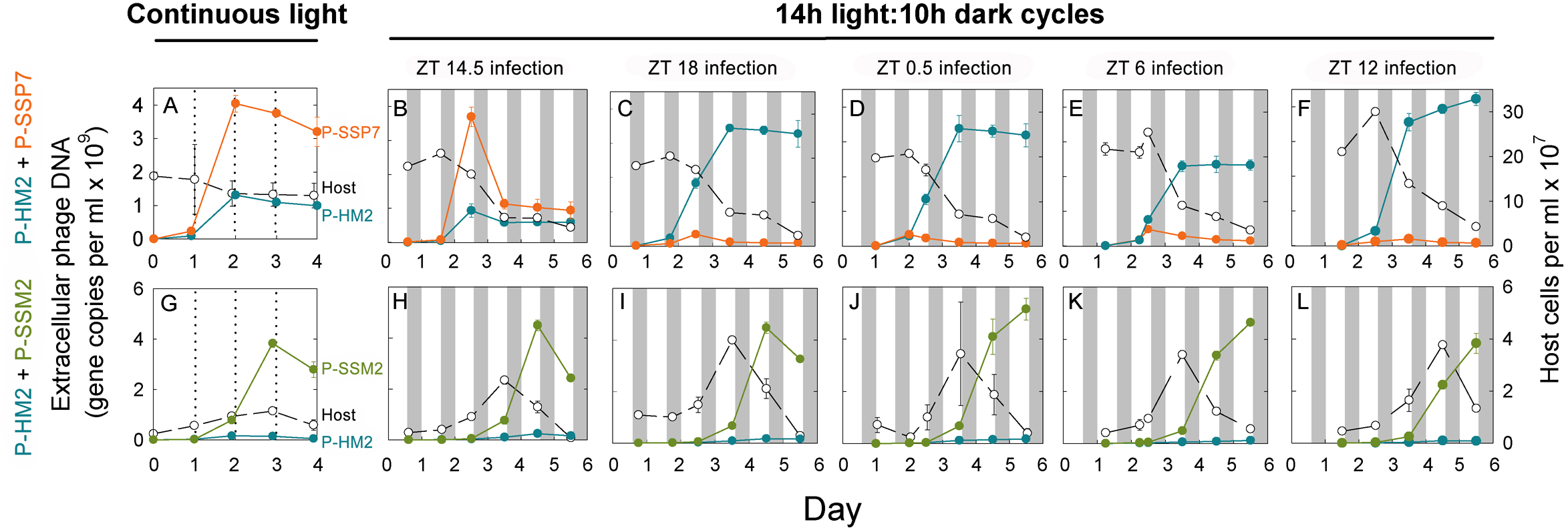
Coinfection of *Prochlorococcus* by different cyanophage pairs. Equal numbers of two cyanophage strains were used to coinfect synchronized *Prochlorococcus* cells at a total phage/host ratio of 0.02. **A**–**F**: *Prochlorococcus* MED4 was coinfected by cyanophages P-HM2 and P-SSP7. **G**–**L**: *Prochlorococcus* NATL2A was coinfected by cyanophages P-HM2 and P-SSM2. Infections were carried out under continuous light (**A** and **G**) or at different times under a 14h light:10h dark cycle (**B** and **H** at ZT 14.5, **C** and **I** at ZT 18, **D** and **J** at ZT 0.5, **E** and **K** at ZT 6, **F** and **L** at ZT 12). Extracellular phages were measured by quantitative PCR with phage specific primers (left y-axis). Host cell numbers were determined by flow cytometry (right y-axis). Error bars represent the standard deviations of two biological replicates.

## Rhythmic variation of cyanophage abundances in the South China Sea and the Western Pacific Ocean

We measured the diel variation of cyanophage abundances in the South China Sea and the Western Pacific Ocean (Figure 4A). During our sampling periods, *Prochlorococcus* was the dominant phytoplankton in both sites and started to divide at around sunset (Figures 4B, D), similar to their growth patterns in other oligotrophic regions (Vaulot et al., 1995; Liu et al., 1997). We carried out plaque assays (Avrani et al., 2011) and found that cyanophages infecting *Prochlorococcus* strains MED4 and MIT9301 were the dominant cyanophages in the South China Sea and the Western Pacific Ocean, respectively (Supplementary Figure 4). During the 36-h sampling period in the South China Sea, the abundance of cyanophages infecting *Prochlorococcus* MED4 increased to a plateau at night and decreased after sunrise (Figure 4B). A similar trend was observed in *Prochlorococcus* MIT9301 phages over 5 days in the Western Pacific Ocean (Figure 4D). Furthermore, the abundances of cyanophages infecting several other *Prochlorococcus* and *Synechococcus* strains were also higher at night than those during the daytime (Supplementary Figure 4). Our results are consistent with the abundance of cyanophages infecting *Synechococcus* WH7803 at 10 m in the Indian Ocean (Clokie et al., 2006). Together, these results indicated that cyanophage abundances in the oceans varied rhythmically under natural light-dark cycles.

**Figure 4.**
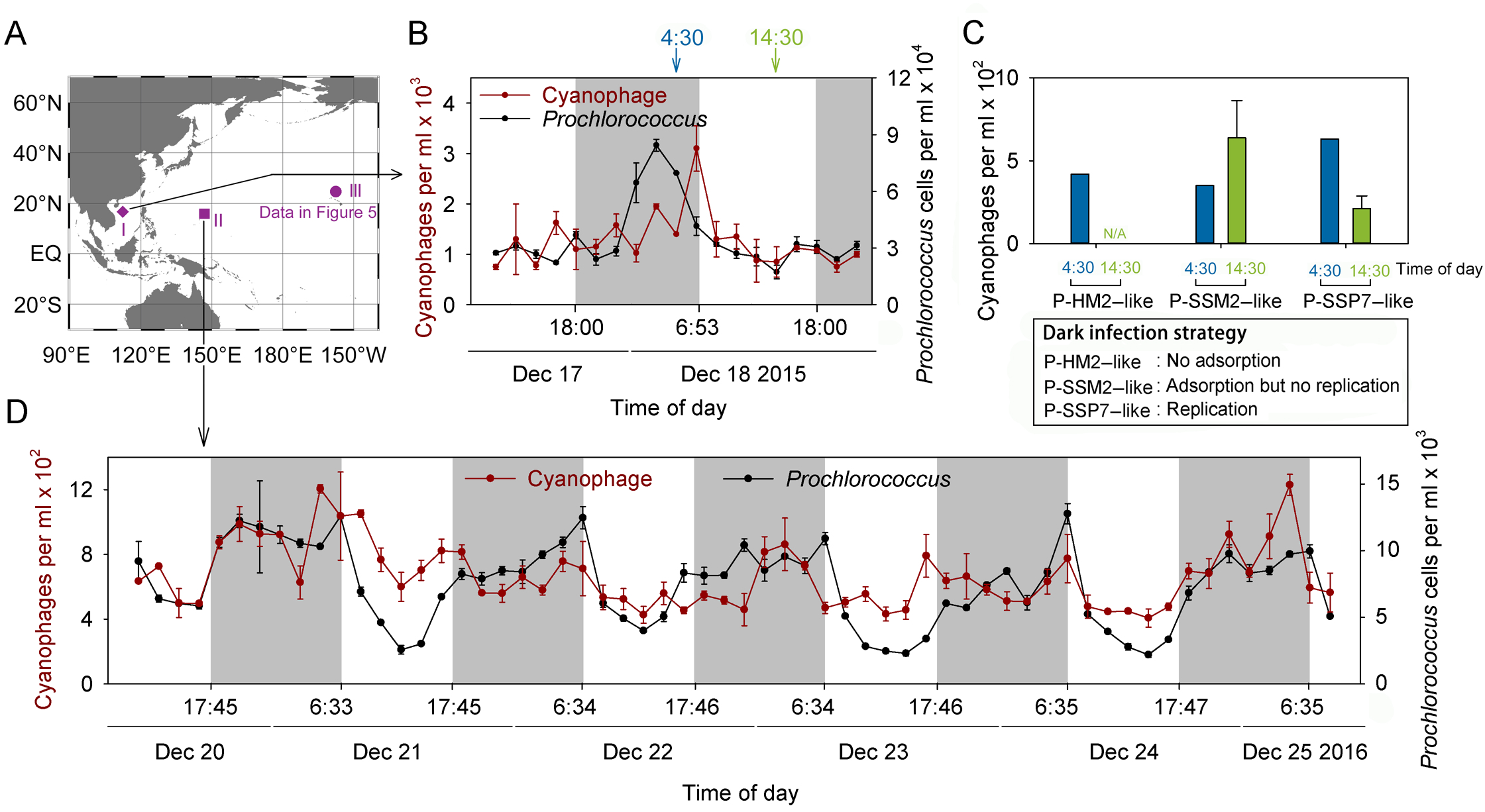
Diel variation of cyanophage abundances in the South China Sea and the Western Pacific Ocean. **A.	**Map showing the sampling sites in the South China Sea (I), the Western Pacific Ocean (II), and the North Pacific Subtropical Gyre (III). The map was generated by Ocean Data View (Schlitzer, 2011). **B**. Surface waters were sampled every 2 h in the South China Sea (16°50′N, 112°19′E). *Prochlorococcus* cells were counted by flow cytometry and plaque assays were carried out to measure the number of cyanophages infecting *Prochlorococcus* MED4. **C**. From *Prochlorococcus* MED4 phages isolated at 4:30 and 14:30 on December 18 2015, the abundances of phages with dark infection strategies similar to P-HM2, P-SSM2, or P-SSP7 were shown. **D**. The numbers of *Prochlorococcus* and cyanophages infecting *Prochlorococcus* MIT9301 in the Western Pacific Ocean (15°42′N, 147°10′E). Error bars represent the standard deviations of two biological replicates.

The decrease of cyanophage abundances after sunrise (Figures 4B, D) could be due to solar radiation, which is the primary factor for virus decay in aquatic environments (Suttle and Chen, 1992; Noble and Fuhrman, 1997). To test whether cyanophages with different dark infection strategies decreased similarly after sunrise, we used *Prochlorococcus* MED4 as the host to isolate 20 phages from the nighttime sample (4:30) and 20 from the daytime sample (14:30) from the South China Sea. After determining their dark infection strategies, we estimated their abundances at 4:30 and 14:30 (Figure 4C). The decrease of P-SSP7–like phages from 4:30 to 14:30 was probably due to solar radiation (Figure 4C). The disappearance of P-HM2–like phages at 14:30 might be caused by both solar radiation and adsorption to the host cells after sunrise (Figure 4C). Interestingly, P-SSM2–like phages increased from 4:30 to 14:30 (Figure 4C). This might be explained by their adsorption to the host cells at night and progeny release in the afternoon, although we cannot exclude the possibility that they are less sensitive to solar radiation. The different trends of three cyanophage groups (Figure 4C) suggested that the dark infection strategy might affect diel variation of cyanophage abundances in natural aquatic environments. We also noticed that the combined abundances of P-HM2–like and P-SSM2–like phages were higher than those of P-SSP7–like phages at 4:30 and 14:30, supporting our hypothesis that cyanophages with infection rhythms have a fitness advantage under light-dark cycles.

To study the diversity of the 40 cyanophages isolated from the South China Sea, we used cyanophage primers to screen them (Sullivan et al., 2008; Sabehi et al., 2012; Dekel-Bird et al., 2015). We found that one cyanophage can be amplified by cyanopodovirus primers and 13 can be amplified by cyanomyovirus primers. While the isolated cyanopodovirus can replicate in the dark, the isolated cyanomyoviruses used three dark infection strategies. Phylogenetic analysis did not cluster cyanomyoviruses according to their dark infection strategies (Supplementary Figure 5), indicating that the dark infection strategy of a cyanophage is not shaped by its phylogeny.

## Diel transcriptional rhythms of cyanophage genes in the North Pacific Subtropical Gyre

To study the diel transcriptional dynamics of cyanophage genes in the oceans, we reanalyzed microbial community transcriptomic data from a previous study conducted over 3 days in the North Pacific Subtropical Gyre (Ottesen et al., 2014). In this dataset, *Prochlorococcus* transcripts were the most abundant, followed by *Pelagibacter* transcripts (Ottesen et al., 2014). Although viral transcripts were much less abundant than their host transcripts, we were able to identify transcripts mapped to phages of *Prochlorococcus*, *Synechococcus*, and *Pelagibacter*, together with several other phage groups (Figure 5A). Since *Synechococcus* is not abundant in this region (Ottesen et al., 2014), we suspected that some *Synechococcus* phages infected *Prochlorococcus*, which has been found previously (Sullivan et al., 2003). We performed harmonic regression analyses following Ottesen et al. (Ottesen et al., 2014) and found that *Prochlorococcus* phages and *Synechococcus* phages showed significant diel rhythm in their total transcript abundances, while other phage groups did not (Figure 5A).

**Figure 5.**
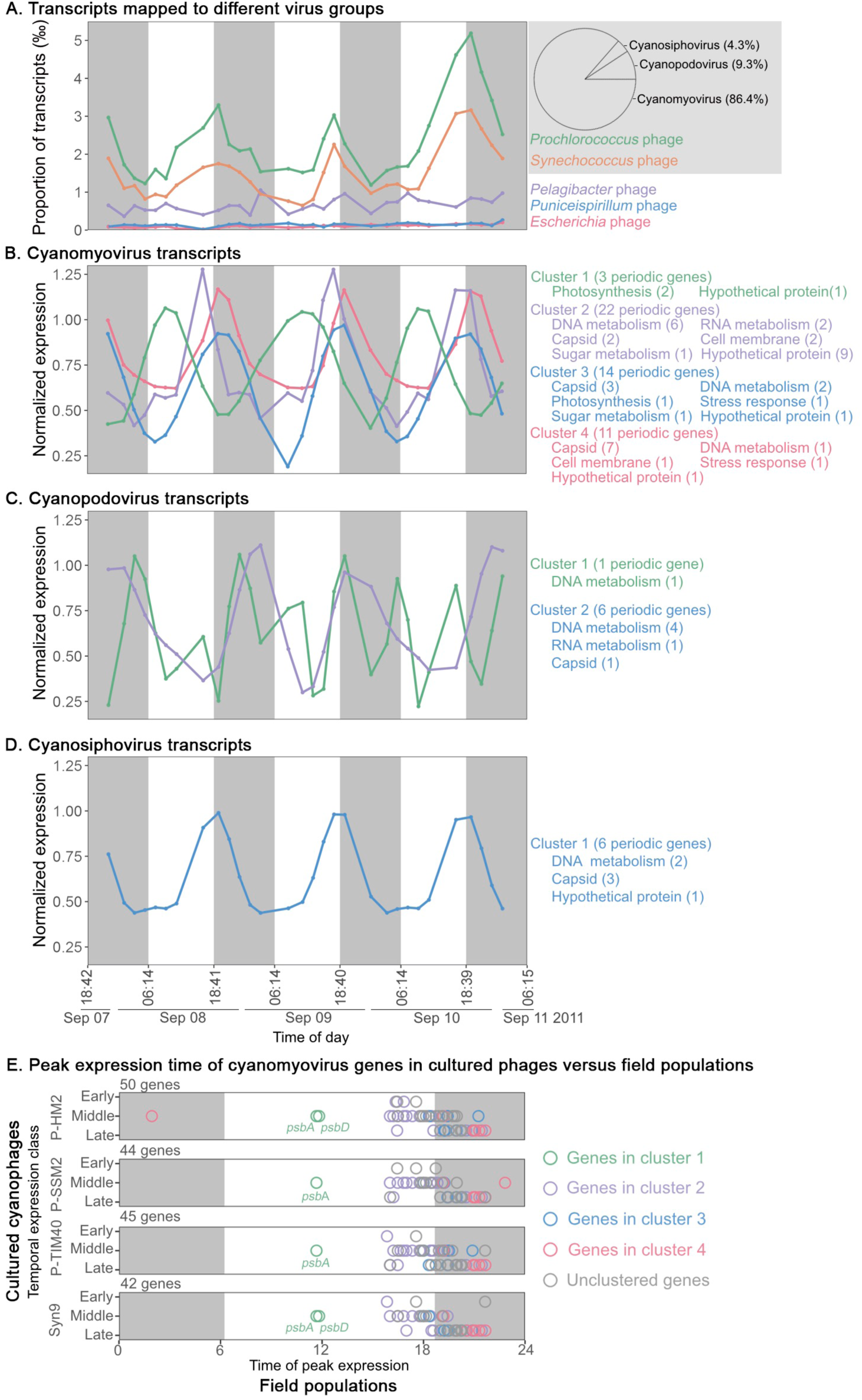
Diel transcript abundances of viral genes in the North Pacific Subtropical Gyre. **A**. Proportion of transcripts mapped to different virus groups at each sampling time point. The pie chart on the right showed the proportion of total cyanophage transcripts (*Prochlorococcus* and *Synechococcus* phages) mapped to cyanomyoviruses, cyanopodoviruses, and cyanosiphoviruses. **B**–**D** Normalized expression levels of periodic genes encoded by cyanomyoviruses (**B**), cyanopodoviruses (**C**), and cyanosiphoviruses (**D**). Each curve showed the average expression level of periodic genes in a cluster, which had similar peak expression times. Twenty-one cyanomyovirus genes cannot be clustered. The expression levels of individual genes were shown in Supplementary Figure 6. **E**. Peak expression time of cyanomyovirus transcripts. The temporal expression class (early, middle, or late) of a gene was assigned based on its ortholog in cyanomyoviruses P-HM2, P-SSM2, P-TIM40, or Syn9. The number of orthologous genes found in each cyanomyovirus genome was shown above the corresponding graph. Different colors indicated the gene clusters shown in **B**. Unclustered genes were indicated by grey color.

Transcripts from *Prochlorococcus* and *Synechococcus* phages were mainly from cyanomyoviruses (86.4%), followed by those from cyanopodoviruses and cyanosiphoviruses (Figure 5A). 71, 7, and 6 periodically expressed genes were identified in cyanomyoviruses, cyanopodoviruses, and cyanosiphoviruses, respectively (Supplementary Figure 6). Many of these genes can be clustered based on their peak expression time (Figures 5B, C, D) and most had peak expression at around sunset, which explained the diel rhythm of total cyanophage transcripts (Figure 5A). The periodically expressed genes were related to DNA metabolism, RNA metabolism, and phage capsid formation (Supplementary Figure 6). For cyanomyoviruses, auxiliary metabolic genes relate to photosynthesis, sugar metabolism, stress response, and cell membrane modification were also found (Supplementary Figure 6), which were thought to redirect host metabolism for cyanophage replication (Breitbart et al., 2007; Thompson et al., 2016).

We compared the temporal expression patterns of cyanomyovirus genes in the field populations with their orthologs in lab cultures. Cyanomyoviruses P-HM2, P-SSM2, P-TIM40, and Syn9 were found to have early, middle, and late temporal gene expression classes (Doron et al., 2015; Lin et al., 2016; Thompson et al., 2016). While most orthologous genes of the four cyanophages are in the same temporal class, some are in different classes (Thompson et al., 2016), probably because of different host cells and growth conditions. Of the 71 periodically expressed cyanomyovirus genes in field populations, 50, 44, 45, and 42 had orthologs in P-HM2, P-SSM2, P-TIM40, and Syn9, respectively. Overall, the early genes of cultured cyanophages seemed to have an earlier peak expression time in the field population and the late genes seemed to have a later peak expression time (Figure 5E). The coordinated expression of early, middle, and late genes in field populations suggested that their infections were synchronized to start in the morning and to lyse the host cells at night, which was consistent with the peak abundances of field cyanophages at night (Figure 4). There were several differences between cyanophage lab cultures and field populations (Figure 5E), which might be due to different genetic backgrounds of their host cells and growth conditions, as has been observed in *Prochlorococcus* transcripts in the field (Ottesen et al., 2014). One example is the expression of photosystem II genes *psbA* and *psbD* in Cluster 1 (Figures 5B, E). They were expressed as middle genes in the lab cultures (Doron et al., 2015; Lin et al., 2016; Thompson et al., 2016) but expressed as early genes in field populations (Figure 5E), which might be regulated to enhance the host light reactions during infection.

## Discussion

Using cultured cyanophages we discovered three dark infection strategies (Figure 1) that are also used by field populations (Figure 4C). Since phage infection of heterotrophic bacteria has not been shown to be affected by light/darkness, these strategies are likely to be shaped by cyanobacteria. On the other hand, cyanophage genes might also play a role since cyanomyoviruses with different dark infection strategies were isolated from the same host strain (Supplementary Figure 5). Future studies are needed to understand the molecular mechanisms underlying the three dark infection strategies.

With limited experimental data, we can only speculate on how the three dark infection strategies affect the fitness of cyanophages. P-HM2 misses its chance to infect host cells at night, however, it may achieve a larger burst size during the daytime when host cells are metabolically more active. On the other hand, P-SSP7 infects host cells at every opportunity, although host cells may not be fully utilized to produce enough progeny phages at night. This strategy may confer P-SSP7 a fitness advantage if cyanophages are competing for a limited number of host cells (Figure 3B). Interestingly, P-SSM2 seems to have the combined advantages of P-HM2 and P-SSP7—it does not miss the chance to be adsorbed to a host cell at night and fully utilizes the host’s resources by replicating only during the daytime. This might be why P-SSM2–like cyanophages are more abundant in the oceans (Figure 4C). With different dark infection strategies, cyanophages may reduce their competition by infecting the host cells at different times of a day. Thus, we propose that the daily light-dark cycle provides cyanophages a temporal niche to promote their coexistence and diversity in aquatic environments.

Diurnal rhythms have been shown for all the three kingdoms of life (Edgar et al., 2012), and our results with lab cultures (Figure 2) and field populations (Figures 4, 5) demonstrated that cyanophages can also exhibit rhythmic infection patterns under light-dark cycles. The peak abundances of cyanophages (Figure 4) and their transcripts (Figure 5) at night in our field data suggested that *Prochlorococcus* infection is synchronized in the oceans, with cell lysis at night. Our results are consistent with the light-dark synchronized mortality of *Prochlorococcus* cells in surface waters, which also reaches a peak at night (Ribalet et al., 2015). Synchronized infection of *Prochlorococcus* by cyanophages may result in a synchronized release of dissolved organic matter to the marine food web, which eventually influences nutrient and energy cycling in the world’s oceans. Rhythmic infection patterns might also be found in viruses of eukaryotic organisms, since it was recently shown that time of infection and the host’s circadian clock affected the outcome of herpes and influenza A virus (Edgar et al., 2016).

## Acknowledgements

This study is supported by grants to Qinglu Zeng from the National Basic Research Program of China (973 Program) (Project number 2013CB955700), the National Natural Science Foundation of China (Project number 41476147), and the Research Grants Council of the Hong Kong Special Administrative Region, China (Project numbers 16144416, 16103414, and 689813).

## Author contributions

Q.Z. and R.L. designed the experiments; R.L. conducted diel infections of cultured cyanophages and measured cyanophage abundances in the field; Y.C. analyzed metatranscriptomic data; R.Z. and N.J. contributed in sampling in the South China Sea and the Western Pacific Ocean; Y.L. did PCR screening of isolated cyanophages; Q.Z. wrote the manuscript with contributions from all authors.

## Methods

### Culture conditions

Axenic *Prochlorococcus* strains were grown in Port Shelter (Hong Kong) seawater–based Pro99 medium (Moore et al., 2002). Batch cultures were incubated at 23°C in continuous light (25 μmol quanta m^−2^ s^−1^) or a 14h light:10h dark cycle (35 μmol quanta m^−2^ s^−1^ in the light period). Cultures were acclimated in the same condition for at least three months before they were used for the infection experiments.

### Flow cytometry and cell cycle analysis

*Prochlorococcus* cells were preserved by mixing 100 µl culture with 2 µl 50% glutaraldehyde to a final concentration of 1% and were stored at –80°C. Cells were enumerated by a flow cytometer (BD FACSCalibur) with the CellQuestPro software. We followed a published protocol to determine the percentage of cells in each cell cycle stage (Zinser et al., 2009). Briefly, *Prochlorococcus* cells were stained with the DNA stain SYBR Green (Invitrogen) and flow cytometry data were analyzed with the ModfitLT software.

### Quantification of cyanophages

In order to set the phage/host ratio for an infection experiment, total phage particles were filtered through a 0.02 µm Whatman Anodisc filter, stained with 5x SYBR gold (Molecular Probes), and counted under an epifluorescence microscope (Chen et al., 2001; Patel et al., 2007). At least five discrete fields on a filter were photographed using the SPOT Advanced Imaging software and fluorescent dots representing phage particles were counted manually.

During infection, extracellular phage DNA was quantified using a quantitative polymerase chain reaction (qPCR) method (Lindell et al., 2007). Briefly, infected *Prochlorococcus* cultures were filtered through 0.2 μm polycarbonate filters in a 96-well filter plate (Pall). Filtrates containing extracellular phage particles were diluted 100-fold in dH_2_O and were then used as templates for qPCR reactions in a 384-well plate. A qPCR reaction contained 4.6 μl template, 0.2 μl forward primer (10 μM), 0.2 μl reverse primer (10 μM), and 5 μl iTaqTM Universal SYBR Green Supermix. The LightCycler 480 Real-Time PCR System (Roche Diagnostics) was used for thermal cycling, which consisted of an initial activation step of 5 min at 95°C, 45 amplification cycles of 10 s at 95°C and 60 s at 60°C, and a melting curve analysis at the end. The number of cyanophages in each well was quantified using a standard curve generated from phage particles that were enumerated by epifluorescence microscopy. The qPCR primers were listed in Supplementary Table 1.

### Cyanophage adsorption in the light and in the dark

*Prochlorococcus* cells were acclimated under continuous light for at least three months. Four tubes of mid-log cultures were mixed with cyanophages at a phage/host ratio of 0.1 in the dark. Two tubes were moved to an incubator with continuous light and two were kept in the dark throughout the whole experiment (in a box wrapped with aluminum foil in the same incubator). At each sampling time point, 100 µL culture was filtered through a 0.2 μm filter. Extracellular phages in the filtrate were quantified by qPCR (cyanophages P-HM1, P-HM2, P-SSP7, P-GSP1, and P-SSM2), or were stained with SYBR gold and counted under an epifluorescence microscope (cyanophages isolated in this study from the South China Sea). Cultures kept in the dark were sampled in a dark room with a dim red light on, which did not affect the adsorption of cyanophages to their host cells (Jia et al., 2010) (Figure 1).

### Infection of synchronized *Prochlorococcus* cells under light-dark cycles

*Prochlorococcus* cells were acclimated under light-dark cycles for at least three months and were synchronized, as determined by flow cytometry (Supplementary Figure 2). Mid-log cells were infected at different times of a light-dark cycle. When *Prochlorococcus* was infected by a single phage strain, the phage/host ratio was 0.02. When *Prochlorococcus* was coinfected by two phage strains, the phage/host ratio of each phage was 0.01 and thus the total phage/host ratio was 0.02.

### Cyanophage isolation from the South China Sea and the Western Pacific Ocean

From 10:30 December 17 2015 to 22:30 December 18 2015, seawater was sampled using a 10 L carboy every 2 h from surface waters off the pier of Yongxing (Woody) Island in the South China Sea (16°50′N, 112°19′E). The local time for sunrise was 6:53 and the time for sunset was 18:00. Water temperature varied between 25.4°C and 26.1°C. From 11:00 December 20 2016 to 9:00 December 25 2016, seawater was sampled using a 10 L carboy every 2 h from surface waters in the Western Pacific Ocean (15°43′N, 147°11′E). The local time for sunrise was 6:36 and the time for sunset was 17:45. Water temperature varied between 27.4°C and 28.0°C. Subsamples for flow cytometry analysis were fixed with glutaraldehyde and stored in liquid nitrogen. Subsamples for cyanophage isolation were passed through 0.2 μm polycarbonate filters and stored in acid-washed glass tubes in the dark at 4°C. We modified a pour plating protocol (Avrani et al., 2011) to carry out plaque assays to count and isolate cyanophages infecting a *Prochlorococcus* strain. Invitrogen UltraPure low melting point agarose (0.37% final concentration) was dissolved in the Pro99 medium supplemented with 1 mM Na_2_SO_3_. The mixture was cooled down to around 27°C, added with *Alteromonas* sp. strain EZ55 (~10^6^ cells), *Prochlorococcus* (~10^8^ cells) and diluted seawater samples, and poured into a 90 mm petri dish. Plates were incubated at 23°C under continuous light (25 μmol quanta m^−2^ s^−1^). The number of plaques was recorded every two days until no new plaque appeared. Well isolated plaques were picked and were used to infect *Prochlorococcus* cells.

### PCR screening and phylogenetic analysis of cyanophages isolated from the South China Sea

Using the 40 cyanophages we isolated from the South China Sea as templates, PCR screenings were done with published primers to amplify signature genes of T4-like (Sullivan et al., 2008), T7-like (Dekel-Bird et al., 2015), and TIM5-like (Sabehi et al., 2012) cyanophages. PCR reactions were performed as previously described (Sullivan et al., 2008; Sabehi et al., 2012; Dekel-Bird et al., 2015) and PCR primers were listed in Supplementary Table 1. After PCR screening, 13 cyanophages can be amplified by T4-like phage primers, 1 can be amplified by T7-like phage primers, and none can be amplified by TIM5-like primers.

After signature genes of cyanophages were sequenced, *g20* amino acid sequences were used to build a phylogenetic tree for cyanomyoviruses. Phylogenetic analyses were accomplished with the Phylogeny.fr website (Dereeper et al., 2008). The distance programme ProtDist/FastDist + BioNJ (Gascuel, 1997) was used for neighbour-joining (NJ) analyses, and the programme PHYLML (Guindon and Gascuel, 2003) was used for the maximum likelihood (ML) analyses. Bootstrap analyses were carried out with 1000 replicate data sets for neighbour-joining (NJ) trees and 400 for maximum likelihood (ML) trees. The final tree topology was implemented with the iTOL:Interactive Tree Of Life website (Letunic and Bork, 2016).

### Metatranscriptomic data analysis

In a previous study (Ottesen et al., 2014), microbial community RNA at 23 m depth in the North Pacific Subtropical Gyre was sequenced from September 7 to 11, 2011. In the original analysis of this study, the authors mainly focused on reads mapped to bacteria and archaea. We downloaded their metatranscriptomic data from NCBI Sequence Read Archive (accession number SRP041215) and analyzed reads mapped to viral genomes. Following published protocols (Ottesen et al., 2013; Ottesen et al., 2014; Aylward et al., 2015), we mapped reads to viral genomes, identified periodically expressed genes, and clustered them by their expression patterns. Below were brief summaries of metatranscriptomic data analysis.

#### Sequence analysis and annotation

Downloaded sequences were quality-filtered as previously described (Aylward et al., 2015). After removing rRNA sequences, putative coding sequences were identified by aligning sequences to an in-house protein database based on RefSeq Archaea, RefSeq Bacteria, RefSeq Protozoa, RefSeq Plant, and RefSeq Viral (RefSeq release 79), together with coding sequences of additional cyanophages (Sullivan et al., 2010; Labrie et al., 2013; Puxty et al., 2015; Crummett et al., 2016). For sequence alignment, LAST (Kielbasa et al., 2011) was used with a bit score cutoff of 50.

Megan (Huson et al., 2007) was used to classify translated coding sequences into different taxonomic groups, based on the best hit of a sequence. Sequences assigned to dsDNA viruses by Megan were divided into several virus groups based on their host taxonomy. If a translated sequence was aligned equally well to different virus groups, or even to both a virus and its host, the DNA sequence was used for taxonomic classification. Ambiguous sequences that cannot be classified into a single group were discarded from further analysis.

For *Prochlorococcus* and *Synechococcus* phages, homologous genes found in multiple phage genomes were grouped into orthologs as previously described (Aylward et al., 2015). Cyanophage orthologs were further grouped into cyanomyoviruses, cyanopodoviruses, and cyanosiphoviruses (Supplementary Table 2).

#### Identification of periodically expressed genes

To identify periodically expressed genes, cyanophage orthologs with more than 30 total read counts were undergone regression tests as previously described (Ottesen et al., 2013). Count data of each ortholog was fitted to the following formula through Poisson log-linear regression: 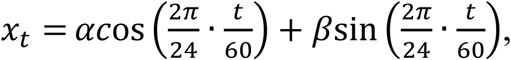 where *t* is the midpoint of the sampling time in minutes. The library offset for regression is the total number of sequences with significant hits (bit score > 50). Chi-square tests and the permutation tests were conducted to assess the significance level of model fit. The chi-square test calculated the *p* values for both variables in the above formula. For each gene, the smaller *p* value generated from chi-square test and the *p* value from the permutation test were taken to measure the significance of model fit. P values from both tests were corrected through FDR controlling procedure. If FDR-corrected *p* values from the chi-square test and the permutation test were both smaller than 0.1, a gene was identified as a periodically expressed gene.

#### Clustering of transcriptional profiles

Periodically expressed genes from cyanomyoviruses, cyanopodoviruses, and cyanosiphoviruses were clustered through an ARMA process using the EM algorithm (Li et al., 2010) as previously described (Ottesen et al., 2013). The read count of each ortholog was scaled by the total count at each time point to calculate its transcript abundance. To cluster orthologs based on their peak expression times, the transcript abundance was normalized by the daily maximum transcript abundance, which was intended to eliminate the effect of the daily amplitude of transcript abundance. The square root and arcsine transform were applied to the relative transcript abundance to stabilize the variance. The formula for data normalization was shown below:

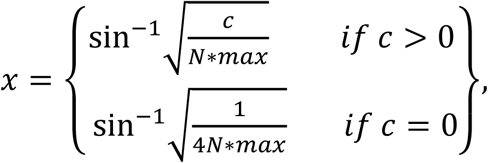

In this formula, *c* represents the count of unique sequences under each ortholog, *N* is the total number of sequences with significant hits at each time point, and *max* is the daily maximum of *c/N*. The optimal model parameters were determined through parameter simulation. Cluster models with different parameter combinations (*P* = 1-2, *Q* = 0-1, *K* = 1-3, *J* = 1-10) were evaluated and compared using the Bayesian Information Criterion. The probability of cluster membership should be larger than 90% to assign a gene to a certain cluster.

